# Somatostatin neurons in the rostral nucleus of the solitary tract are functionally heterogeneous

**DOI:** 10.64898/2026.04.21.719923

**Authors:** B. Kalyanasundar, Charlotte Klimovich, Susan Travers

## Abstract

The rostral nucleus of the solitary tract (rNST) is the initial central site for taste processing. This nucleus has a complex circuitry and multiple cell types with different response properties, connectivity, and morphology (Travers and Travers 2018). However, unlike its visceral counterpart, the caudal NST, neurochemical phenotypes in rNST are poorly defined. Recent studies have begun to probe this gap. Based on fiber photometry, optogenetics, and cell-type specific deletion. For example, one group proposed that somatostatin (SST) rNST neurons, neither calbindin or dynorphin cells, responded specifically to bitter stimuli and that these neurons were necessary for suppression of quinine-induced licking (Jin, Fishman et al. 2021) (Zhang, Jin et al. 2019). The present study employed in situ hybridization, optotagging, and chemogenetic suppression in male and female mice to demonstrate that SST neuron function is more complex. Although most SST neurons responded optimally to bitter stimuli, many others were activated by different qualities and some non-SST neurons responded to bitter tastants. Moreover, roughly equal proportions of SST neurons expressed excitatory (VGLUT2) or inhibitory (VGAT) markers. Suppressing SST neural activity with DREADDS enhanced licking to both quinine and sucrose suggesting that neural activity elicited by the aversive bitter stimulus was suppressed whereas licking elicited by the sweet, preferred stimulus was increased. We hypothesize that these effects arise from suppressing excitatory quinine-responsive SST neurons but that a separate population of inhibitory SST neurons synapse on sucrose-responsive cells.

**Significance Statement:** Recent studies have revealed molecular heterogeneity of gustatory system neurons. However, it is unclear whether molecularly-distinct cells are associated with specific roles. The current study investigated somatostatin (SST) neurons in rNST, the first central hub for taste processing. Well over half were inhibitory, expressing VGAT, but a substantial proportion were excitatory, expressing VGLUT2. A narrow majority responded optimally to the bitter quality and none to NaCl, but other SST cells responded most vigorously to sweet, umami, or sour stimuli. Subsets of neurons not expressing SST responded best to each quality, including bitter. Suppressing activity in SST neurons dampened behavioral avoidance to quinine but enhanced consummatory responses to sucrose. Thus, SST rNST neurons exhibited varied functional characteristics but also clear distinctiveness.

## INTRODUCTION

The rostral nucleus of the solitary tract (rNST) is the initial central hub for taste processing. This nucleus includes subdivisions with morphologically distinct cell types that have preferential afferent and efferent connectivity (Whitehead 1988, Halsell, Travers et al. 1996, Ganchrow, Ganchrow et al. 2014, Jin, Fishman et al. 2021). Moreover, the rNST includes ample populations of excitatory and inhibitory neurons (Wang and Bradley 2010, Boxwell, Yanagawa et al. 2013). Other characteristics indicative of complexity are responses that exhibit multisensory integration (Travers and Norgren 1995, Lemon 2017), plasticity as a function of metabolic and dietary variables (Giza and Scott 1983, Nakamura and Norgren 1995, Mai, Ramos et al. 2025) and modulation by behavioral context (Deng, Xiao et al. 2021, Pilato, O’Connell et al. 2024). The representation of gustatory quality in single cells has been a subject of much debate, with polar views emphasizing broad or narrow tuning, and the lack or presence of chemosensitive cell types (reviewed in (Spector and Travers 2005, Ohla, Yoshida et al. 2019, Travers and Spector 2021). Recent investigations cast a new light on this question by evaluating taste responses in phenotypically distinct cell populations made possible by gaining access to genetically defined neuron types (reviewed in (Nakajima 2024). Using fiber photometry, Ca^++^ imaging, optogenetics, and cell-type specific deletion, one group proposed that rNST dynorphin-expressing neurons are selectively responsive to sour stimuli whereas calbindin and somatostatin (SST) neurons are responsive to sweet and bitter tastants, respectively (Zhang, Jin et al. 2019, Jin, Fishman et al. 2021). This may be an oversimplified view, especially for SST neurons. Of significance, the caudal NST also contains a substantial group of SST cells. Some are excitatory and express a marker for glutamate, VGLUT2. However, a majority are inhibitory, express markers for GABA (GAD65, VGAT) and exert inhibitory physiological effects (Bellusci, Garcia DuBar et al. 2022). With this in mind, we hypothesized that SST rNST neurons may also be heterogeneous. The present manuscript identified the excitatory/inhibitory phenotype of SST rNST neurons using in situ hybridization. We then determined the chemosensitivity of optopgenetically-identified SST cells and probed their behavioral function using DREADDS to suppress neural activity during brief-access gustatory testing

## MATERIALS and METHODS

The Ohio State University IACUC approved all procedures involving animals (Protocol 2009A0220).

### IN SITU HYBRIDIZATION

We used in situ hybridization to determine the excitatory and inhibitory phenotypes of SST neurons in the rNST. Six mice (3M, 3F, 63 ± 5 days old) were deeply anesthetized with ketamine (150 mg/kg) and xylazine (20 mg/kg), I.P. and then perfused through the left ventricle with phosphate buffered saline (PBS) followed by 4% paraformaldehyde. Brains were extracted, post-fixed and cryoprotected (4% paraformaldehyde/30% sucrose) in the refrigerator overnight. Subsequently, 15 µm sections of the medulla were cut on a cryostat and collected into cryoprotectant solution for storage at -20°C until mounting and processing.

For the in-situ procedure, we chose sections from the caudal, middle, and rostral thirds of rNST. An additional section, usually from the caudal NST, was processed as a negative control. We washed sections in DEPC-PBS before mounting onto Gold Seal Ultrastick Adhesion slides where the remaining in situ processing took place. In situ processing used the RNAscope Multiplex Fluorescent Reagent Kit v2 and followed the suggested protocol from the manufacturer with minor adjustments. We used probes to detect mRNA for SST (404631, Mm-Sst), VGLUT2 (Mm-Slc17a6, #319171) VGAT (319191, Mm-Slc32a1) and a probe against a bacterial molecule (DapB of Bacillus subtilis strain) as the negative control. No staining was detected in these negative control sections. In situ staining was carried out with 3 probes simultaneously: VGLUT2, VGAT and SST with labeling using Akoya Opal dyes (Opal 520,#FP1487001KT; Opal 570, #FP1488001KT; Opal 690, # FP1497001KT).

To facilitate drawing NST borders, we acquired 10X photomicrographs using a Nikon E600 microscope that included a darkfield channel. Subsequently, for analysis, we took four 20X confocal stacks using an Olympus FV3000 microscope (resolution = 0.621 µm in the XY axis, 2µm separation in the Z plane, excitation wavelengths = DAPI (461nm), Opal 520 (488nm), Opal 570 (561nm), and Opal 690 (640nm)). The four stacks were assembled into a composite image with the stitching module in Image J (Preibisch, Saalfeld et al. 2009).

Using the composite images, cells were counted manually using the Image J Cell Counter module. SST-labeled cells were identified and then classified as co-stained for VGLUT2 and/or VGAT. To equalize the appearance of the images on the different channels, initial cell counts were assessed using the grayscale display. As detailed below, a very small proportion of neurons were clearly triple-labeled. However, about 5% of SST neurons stained primarily for VGAT *or* VGLUT had minimal though detectable expression of the other transporter. To simplify analysis, these neurons were classified as double-labeled for the most strongly stained transporter.

To define the distribution of labeled cells in the coronal plane, we divided the nucleus into mediolateral thirds and dorsal and ventral halves to yield “subfields” (King, Travers et al. 1999, Travers, Herman et al. 2007): medial dorsal (MedD), mid dorsal (MidD) lateral dorsal (LatD), medial ventral (MedV), mid ventral (MidV), lateral ventral (LatV). To account for differences in the sizes of the nucleus at different AP levels and different subfields, raw counts were converted to densities when analyzing the anatomical distributions.

### NEUROPHYSIOLOGY

#### Mice

Mice (N=19; 10 females, 9 males) were crossed between homozygous Sst-IRES-Cre (Jackson Labs #028864) and floxed ChR2/EYFP (Jackson Labs #024109) mice. Subjects were adults and ranged in age from 51-347 days (mean = 225.0 ± 102.3 days, SD).

#### Surgical Preparation for Recording

To anesthetize mice for surgery, animals were sedated with isoflurane (2%) in a chamber and subsequently injected with urethane (1g/kg, I.P.). Throughout surgery and recording, the level of anesthesia provided by this single dose of urethane was supplemented with isoflurane (0.5-1% in 450-600 mmHg oxygen) titrated to maintain an areflexive state To view the mouth clearly and create conditions optimal for whole mouth stimulation, we passed sutures through the labial commissures and another through the lingual mucosa anterior to the foliate papillae to retract the lips and pull the tongue laterally and anteriorly. In addition, we severed the hypoglossal nerves and inserted a tracheal cannula (Breza and Travers 2016, Kalyanasundar, Blonde et al. 2020, Kalyanasundar, Blonde et al. 2023). After placing the animal in a stereotaxic device, an incision was made through the skin over the skull dorsum prior to using a drill and rongeurs to remove the interparietal plate over rNST. To provide unfettered oral access, we glued a small stainless-steel bolt to the skull between bregma and lambda to stabilize the head and removed the stereotaxic mouthpiece.

#### Taste Stimulation

Gustatory stimuli were made with reagent-grade chemicals (Fisher or Sigma) diluted in artificial saliva (AS, in mM: 22 KCl; 15 NaCl, 0.3 CaCl_2_, 0.6 MgCl_2_) (Breza and Travers 2016). Taste stimuli (in mM) were representatives of standard taste qualities: “sweet”, sucrose (SUC, 600), “umami”, a cocktail of MSG (100), IMP (2.5), and amiloride (0.1) (MSGai), “salty”, NaCl (100), “sour”, citric acid (CIT, 60), “bitter” (BIT) a mixture of cycloheximide (0.01) and quinine (2.7). These intensities were mid-range but at the higher end of those in our previous studies (Kalyanasundar, Blonde et al. 2020, Travers, Kalyanasundar et al. 2022, Kalyanasundar, Blonde et al. 2023) and similar to those employed by Jin and colleagues (Jin, Fishman et al. 2021). We used these potent concentrations to increase breadth of tuning (Wu, Dvoryanchikov et al. 2015) and thus better reveal the full sensitivities of individual neurons. Stimuli were delivered from pressurized glass reservoirs connected via polyethylene tubes to a manifold (Warner Instruments) attached to a glass tube (1.0-1.2 mm) positioned to stimulate the entire mouth. We also used a mixture of stimuli associated with different taste qualities primarily as a search stimulus (in mM: sucrose (300), NaCl (100), citric acid (10), and cycloheximide (0.01)). Stimuli were presented at room temperature at a flow rate of 0.3 ml/s. Tastants were applied for 10 s, preceded and followed by AS (10 s, and 20 s, respectively). We separated successive taste stimulations by 1 min or more.

### Neural Recording and Testing Protocol

#### Search Tracks

Experiments commenced by locating gustatory responses in NST, beginning at ∼2.5mm caudal to lambda and 1.0mm lateral to the midline. We recorded neural activity with epoxylite-coated tungsten microelectrodes ∼2-3 mΩ (Frederick Haer Inc. or World Precision Instruments), amplified and filtered the neural signals (10,000X; 600-10K; Alpha Omega, MCPplus), visualized on an oscilloscope and computer display (AD hardware and software, Cambridge Electronics Design [CED], Spike 2), and monitored on a Grass AM8 audiomonitor. Because there is an orderly representation of proprioceptive, tactile, and taste responses in the region (Travers and Norgren 1995), we probed for responses to depressing the jaw, innocuous oral tactile stimulation and tastants to narrow in on the gustatory-dominant zone.

#### Single-unit recording

##### Identifying Taste and Brain-light Responsive Cells

After locating gustatory NST, we switched to recording using an “optrode”, a tungsten electrode (∼2-3 mΩ) glued to a 100 µm, 0.22 nA optical fiber (Thor Labs UM22-100). The optical fiber terminated < 1 mm dorsal to the electrode tip (mean distance = 618 µm; range 450-950 µm, N = 19 measurements). To search for single units, we used taste stimuli and directed blue light pulses (470 nm, 5 ms, 1-10 Hz) to the brainstem via the optical fiber (referred to as light_br_). Light pulses were generated by a laser (Laserglow, LRD-0470-PFFD-00100-05) controlled by CED software and hardware. Prior to recording, we measured the mean light intensity of each of 5 consecutive 5 ms pulses with a Thor PM 100D power meter and averaged across the train. Pulses for the 19 optrodes used in these experiments were 7.9 ± 0.4 mW. The light_br_ is expected to activate both local SST neurons as well as fibers originating from SST neurons outside NST, e.g., the central nucleus of the amygdala (Saha, Henderson et al. 2002, Bartonjo and Lundy 2020, Jin, Fishman et al. 2021).

##### Testing

When a neuron was located that responded to taste or light_br_, further tests were performed. To define a neuron’s gustatory response profile, neurons were systematically tested with the 5 standard stimuli, presented in varied order. Cells that responded to light_br_ in an excitatory fashion were scrutinized further using a 10 Hz/10 s train of blue light pulses and/or a series of 20 pulses at 1, 4, 10, 20 and 50 Hz. Those with short-latency time-locked responses likely expressed ChR2 and SST and thus taste-responsive neurons with these characteristics were classified as SST_pos_. Taste-responsive cells that did <u>not</u> respond in an excitatory, time-locked fashion to light were unlikely to express SST (SST_neg_). However, we were also interested in whether activating nearby SST neurons or fibers modulated taste activity in the recorded cell, and so we tested SST_neg_ cells with gustatory stimuli with and without concurrent light_br_ (10 Hz) stimulation. Stimulations were repeated when possible.

### Neurophysiological Data and Statistical Analysis

#### Basic Measures

##### Taste and Optogenetic Responses

Taste responses were quantified as evoked activity adjusted by a comparable period without taste stimulation using the pre-stimulus AS period to control for any tactile or thermal responses. The pre-stimulus period was calculated separately for control stimulations and those with concurrent light_br_, since activating the SST network sometimes changed spontaneous rate. The mean of repeated trials was the response measure. For each neuron, the mean and standard deviation (SD) of the spontaneous and pre-rinse AS activity were calculated across trials to derive a response criterion. The response criterion was a net evoked firing rate ≥ 1Hz (i.e., at least 10 spikes for 10 s) and 2.5 X the SD of the mean response to AS (Geran and Travers 2006, Kalyanasundar, Blonde et al. 2020); Nishijo, 1991 #411}. However, two neurons that did not strictly meet this criterion were included. One was a bitter-responsive neuron with a clear but delayed response and another a sucrose-responsive neuron that responded with only 0.5 spikes/s, but consistently across 2 trials. Because response rates varied widely across neurons, we derived a second measure, which was net response to a given stimulus divided by the maximum response of that neuron.

##### Brain Light Responses

To analyze responses to light_br_, we triggered off the onset of the stimulus pulse and set a 10 ms search window for detecting a spike. Measures included latency, jitter (SD of latency) and the proportion of trials evoking a spike for each frequency. These measures were averaged across any repeated trials.

#### Additional Analyses

Statistical analyses and graphs were prepared in Systat (v13), Excel (2016), and GraphPad Prism (v9.1.1-10.6). Comparisons between taste responses for SST_pos_ and SST_neg_ neurons were performed using ANOVA. For some analyses, neurons were split into groups based on their optimal responsiveness to representatives of different stimulus qualities: sweet-umami, salty, sour, and bitter. These groups were identical to groups obtained using hierarchical cluster analysis (Pearson’s r; average amalgamation schedule, not shown). Post-hoc t-tests to assess differences in responses for individual stimuli were Bonferroni-adjusted. Comparisons between taste responses under control conditions and during brain light stimulation (light_br_) were compared using repeated measures or mixed ANOVAs. The criterion for statistical significance was set at P<.05.

#### Reconstruction of Recording Sites

##### Marking recording sites

In selected instances, recording sites were marked with an electrolytic lesion produced by passing current (3–8 µA for 3-10 s) at the recording site. In other cases, we made lesions in the reticular formation or vestibular nucleus ventral or dorsal to the cell at a stereotaxically measured distance. At the end of recording, the animal was injected with a lethal dose of anesthesia (80 mg/kg ketamine and 100 mg/kg xylazine) and perfused through the left ventricle with phosphate-buffered saline (PBS), and 4% paraformaldehyde (in 0.1 M phosphate buffer) containing 1.4% L-lysine acetate and 0.2% sodium metaperiodate (McLean and Nakane 1974). The brain was fixed overnight in 20% sucrose paraformaldehyde, blocked in the coronal plane, and stored in sucrose-phosphate buffer prior to sectioning.

##### Histology

Coronal sections of the medulla (40-50 µm) were cut, mounted and stained immediately or stored at –20°C in cryoprotectant until processing. In all but 3 cases, sections were stained with cresyl violet. Reconstructing recording sites in the dorsal-ventral axis was not practical because lesion size was relatively large compared to the thin nucleus. However, for neurons where lesions were made at the recording site or on the same track (N=13), we reconstructed locations as a proportion of their distance from the caudal to the rostral pole and medial to the lateral border of rNST. All neurons were in rNST and recording sites spanned much of the distance in the anterior-posterior (25-93%, mean = 66 ± .06%) and medial-lateral 33-76%, mean = 53%) planes. There was no significant difference in location as a function of whether neurons were SST_pos_ or SST_neg_ or chemosensitivity (ANOVA: anterior-posterior: SST status, P= .989, chemosensitivity, P=.392, interaction, P=.239; medial lateral: SST status, P= .212, chemosensitivity, P=.941, interaction, P=.196).

### BEHAVIORAL STUDIES WITH DREADDs

#### Mice

To study effects of suppressing activity in rNST SST neurons on taste-driven behavior, we used mice that expressed the inhibitory DREADD receptor, hM4Di(Gi) (Urban and Roth 2015) and the fluorescent protein, mCherry or mCherry only, in rNST SST neurons following viral injections (Jackson Labs #028864).

#### Viral Injections

Mice were anesthetized with a cocktail of xylazine and ketamine (2.5 and 25 mg/ml, 0.06 ml/20g bw, I.P.) and maintained with isoflurane (0.5-1%). We disinfected the scalp with alternating swipes of 70% ETOH and iodine, made a midline incision, drilled a small hole through the skull over rNST and removed the dura. We made bilateral 50 nL pressure injections through glass pipettes (tip size = 40-50 µm) using a General Valve Picospritzer. Volume was monitored by observing the meniscus through a microscope equipped with a micrometer. rNST injection coordinates were determined by locating taste-responsive activity with a search electrode and subsequently positioning an injection pipette filled with the virus at the same coordinates. Neural activity through the injection pipette was monitored for assuring accurate depth. Following injection, the pipette was left in place for about 10 min. Mice in the inhibitory DREADD (“Gi”) group were injected with pAAV2-hSyn-DIO-hM4D(Gi)-mCherry, ADDGENE #44362, titre: 1.5 ×10^13^ GC/ml); mice in the mCherry viral control group (“mC”) received pAAV2-hSyn-DIO-mCherry (ADDGENE #50459, titre: 2.6×10E^13^ GC/ml). Mice were assigned to different squads; G_i_/mC - 2 squads; mC - 1 squad).

#### Experimental Design: Behavioral Training and Testing

Training and testing took place in a standard brief-access testing apparatus (Davis MS160-Mouse; DiLog Instruments, Tallahassee, FL, “Davis Rig”), with stimulus presentation and timing under computer control. Mice licked solutions from bottles fitted with sipper tubes that automatically moved into position. Bottle access was controlled with a shutter. The experiment proceeded in three blocks: (1) training, (2) quinine testing, (3) sucrose testing. We tested quinine prior to sucrose because previous observations suggested that mice were reluctant to sample this aversive stimulus after first experiencing sucrose. Stimuli were reagent grade and dissolved in distilled water: quinine (QUI, µM: 30, 100, 1000, 3000), sucrose (SUC, mM: 30, 100, 300, 600, 1000). Training and quinine blocks were performed under ∼23-h water deprivation and occurred daily (Kalyanasundar, Klimovich et al. 2022, Travers, Kalyanasundar et al. 2022). When evaluating sucrose, mice had free access to water but were tested under 23.5-h food deprivation. After the test session, food was returned to their cages until the following day when it was removed again in preparation for the next test. Thus sucrose testing took place on alternate days (Inui-Yamamoto, Blonde et al. 2020, Kalyanasundar, Klimovich et al. 2022, Schier, Inui-Yamamoto et al. 2019, Travers, Kalyanasundar et al. 2022).

Training began ∼two weeks (mean = 2.4 ± 0.5 SD) following the viral injection. During the first two training days, mice were placed in the rig for 30 minutes with the shutter open and a single bottle of distilled water available. On day 3, mice were permitted water access in discrete trials: a bottle moved into position, the shutter opened and remained open for 5 s after the first lick and then closed for 10 s before opening again. There was no restriction on how many trials mice could initiate in the 30-minute period. Days 4 and 5 were identical to day 3, except that we gave I.P. injections of CNO or saline (counterbalanced) prior to the session to adapt animals to the injection procedure and to test for any untoward effects of CNO (we did not observe any). This same 5-s trial structure was used for testing taste stimuli. We tested quinine and sucrose following counterbalanced I.P. injections with saline (∼0.1ml/10g bw, I.P.) and CNO (∼0.1ml/10g bw, I.P.) Altogether, there were 4 sessions for each stimulus (two with saline; two with CNO). Following the four sessions with QUI, we interposed a single SUC session in water-deprived mice to encourage sampling and then proceeded with the four additional sucrose sessions following saline or CNO injections. Following the last test period, mice were perfused as in the neurophysiological experiments and brains cut into 50 µm sections. Slices were mounted onto slides and fluorescent photomicrographs taken of NST, including the injection site. Except for three cases, injections were centered in the rNST bilaterally. In two other cases, injections were unilateral and in a third, no virus was visible. These three cases were excluded from analysis.

### Behavioral and Statistical Analysis

Responses to QUI were quantified as the ratio of the number of licks to QUI relative to the average number of licks to water for a given test day. For SUC, we used a lick score where the number of licks for a given trial was normalized by subtracting the average number of water licks in the session (Schier, Inui-Yamamoto et al. 2019, Travers, Kalyanasundar et al. 2022, Ascencio Gutierrez, Martin et al. 2024, Kalyanasundar, Harley et al. 2025). Responses were averaged across all trials for both sessions with the same stimulus (SUC or QUI) and condition (CNO or saline). [In one SUC session, we noted a consistent dip in licking for 300 mM SUC for all mice tested, inferred that the bottle was not functioning properly, and so dropped that stimulus from analysis on that day.] For the QUI tests, there were 2 mice for which we could use only 1 session for one or both conditions due to inadequate sampling or procedural errors. Analyses were performed using repeated measures ANOVAs with concentration and drug state as the independent variables. These were followed by Bonferroni-adjusted paired t-tests for each concentration. Statistics were performed in Systat (v13), Excel (2016), and GraphPad Prism (v10.4.1). The significance level was set at *p* < .05.

For all experiments, details of statistical analysis are typically presented in figure captions. Errors are S.E.M. unless otherwise noted.

## RESULTS

### Phenotypes of rNST SST neurons

Triple in-situ hybridization for SST, VGAT, and VGLUT2 revealed substantial populations of neurons co-expressing SST with VGAT or VGLUT2 but few expressing only the peptide (FIGURE 1). Across 1,150 SST cells analyzed from 6 mice, Across the six mice and 1,150 SST cells analyzed, neurons expressing SST and VGAT comprised 61.3 ±1.5%, SST and VGLUT 35.5% ± 3.6% of the population. The remainder (2.3 ± 0.2%) expressed only SST or were triple-labeled (1.0 ± 0.04%). Thus, well over half of the SST neurons were inhibitory. Despite the large numbers of both excitatory and inhibitory SST neurons, there was an uneven topographical distribution of the two types of SST neurons in the nucleus (FIGURE 2). In the coronal plane, the densities of SST/VGAT neurons were higher than SST/VGLUT laterally and ventrally. Specifically, inhibitory SST neurons were significantly denser in the LatD, MedV and LatV subfields, but similar elsewhere. Across the anterior-posterior axis, the density of SST/VGAT neurons significantly exceeded that of SST/VGLUT cells at the caudal and middle levels but were similar at the rostral level.

**Figure 1.**
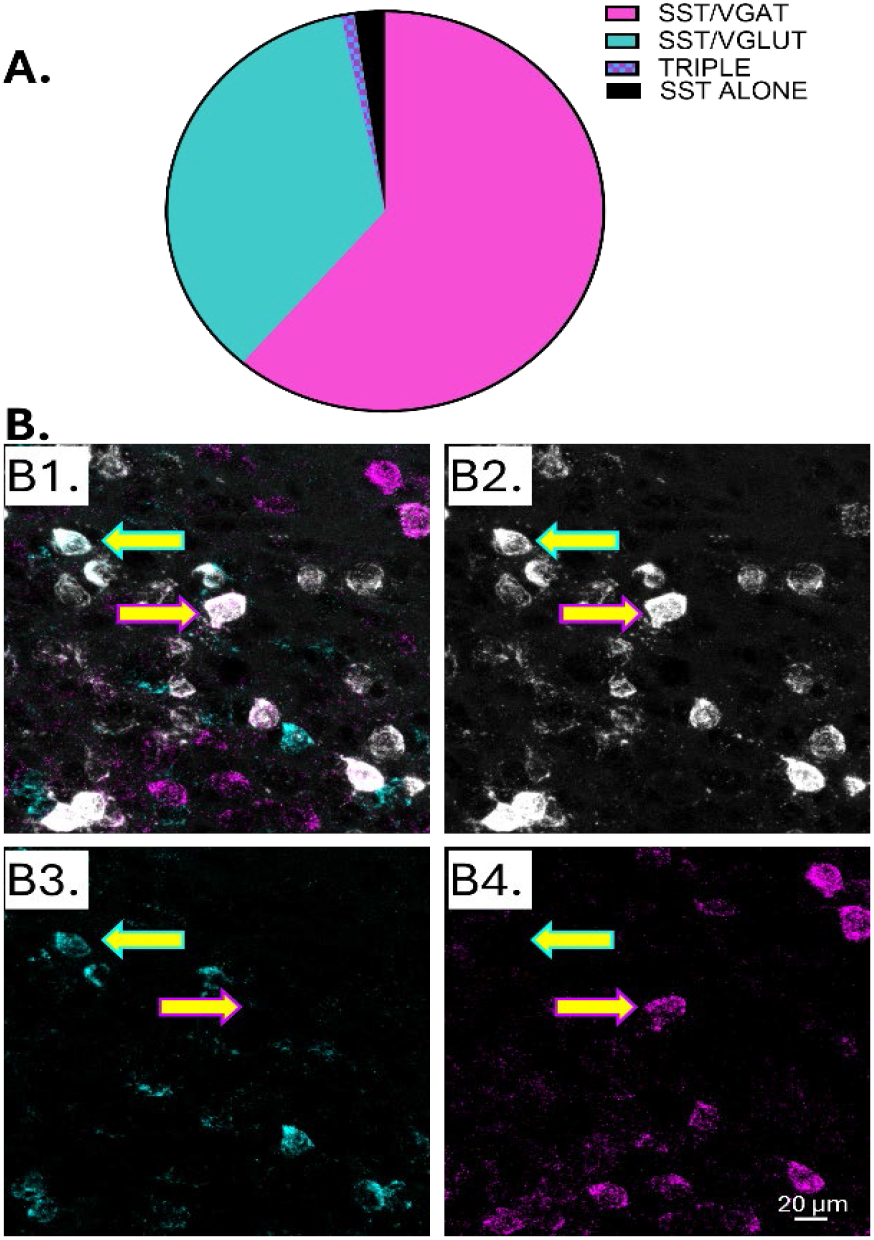
**A**. Proportions of neurons labeled for SST alone, SST and VGLUT, SST and VGAT, or triple labeled. **B**. Confocal photomicrographs showing in situ hybridization for SST (white), VGAT (magenta), and VGLUT2 (cyan). **B1:** overlay, **B2**: SST, **B3:** VGLUT, **B4:** VGAT. Images are maximum intensity projections of nine 1 um planes in the z axis. Brightness of the single channel images was enhanced for clarity.

**Figure 2.**
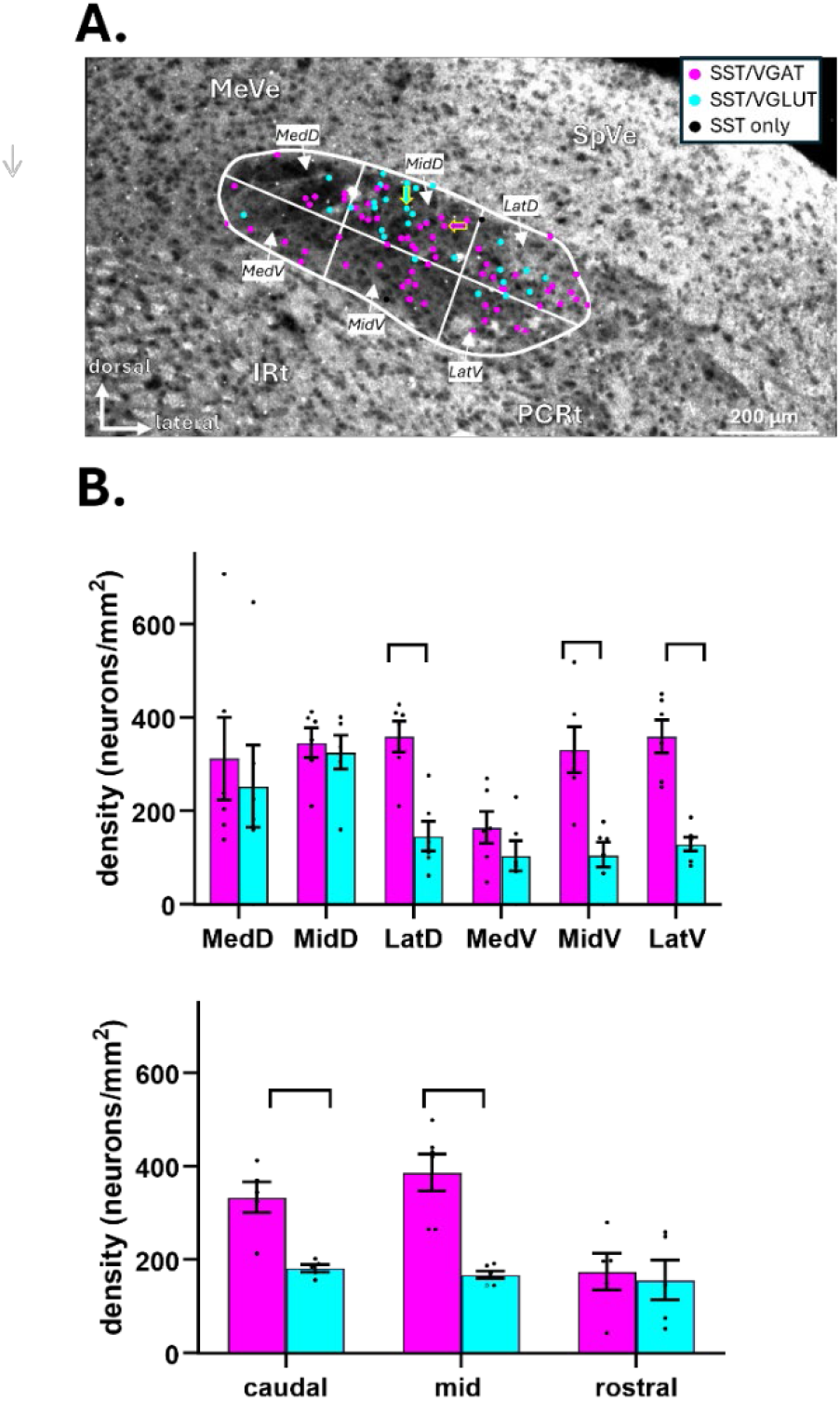
**A**. Plot of classified neurons at a middle rNST level superimposed on a darkfield image. Outline shows the approximate border of the nucleus, and the grid shows divisions into subfields: from medial to lateral, dorsal tier: MedD, MidD, LatD; ventral tier: MedV, MidV, LatV. The arrows point to the same neurons indicated by the arrows in the photomicrograph in Figure 1. Abbreviations: Irt-intermediate reticular nucleus, MeVe: medial vestibular nucleus, PCRt-parvicelluar reticular nucleus. **B**. Densities of the different types of SST neurons across subfields and AP levels. A repeated-measures ANOVA for subfields yielded significant main effects for neuron type (*p* = .001) and subfield (*p* = .004) and an interaction (.015). ANOVA for AP level likewise revealed main effects for both variables (type, *p* = .001; level, *p* = .005) and an interaction (*p* = .003). Brackets above the bars indicate significant differences (*p* < .05) between densities of SST/VGAT and SST/VGLUT2 neurons for a given area based on Bonferronni-adjusted paired T-tests. Individual data points (N =5-6) are indicated.

### Neurophysiological Results

#### Chemosensitivity of SST_pos_ and SST_neg_ neurons

We recorded from 31 taste-responsive rNST neurons in 20 mice. In 14 mice, 432 nm light_br_ evoked a time-locked action potential and were thus considered “optotagged”. Each optotagged cell responded at short latency and reliably; i.e., on at least 80% of pulses delivered at 1 Hz stimulation. In fact, most (10) responded to light_br_ on at least 95% of trials. We tested twelve of the optotagged neurons with the full frequency series (FIGURE 3). These cells maintained short latency and low jitter across 1-100 Hz, although responding was less faithful at the higher frequencies. The 14 optotagged neurons were presumed to be SST_pos_; the remaining 17 cells lacking an excitatory, light-driven response were classified as SST_neg_. In 11 mice, we recorded from only one cell, but multiple cells were studied in the other subjects. In most mice (6/9) where multiple cells were recorded, both SST_pos_ and SST_neg_ cells were encountered, suggesting that types were not dependent on technical variables in a given preparation. The chemosensitivities of SST_pos_ and SST_neg_ neurons overlapped considerably but some differences were observed.

**Figure 3.**
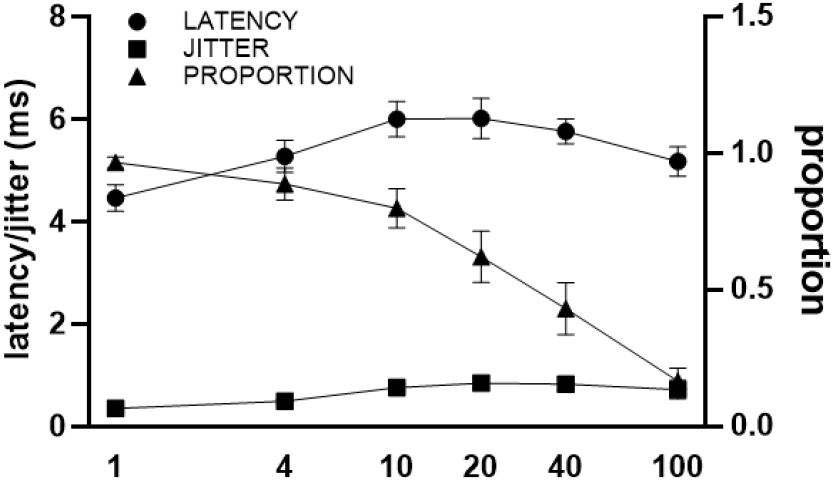
Latency, jitter, and proportion (mean ± S.E.M.) of light-evoked action potentials for light_br_-driven neurons tested with the full frequency range (N = 12). The proportion was nearly unity at 1 Hz but systematically declined, especially at frequencies above 10 Hz. However, the mean latency (4.5-5.2 ms) and jitter (0.35-0.72 ms) of the light_br_-elicited responses was short and relatively stable across frequencies.

Both SST_pos_ and SST_neg_ neurons were responsive to all five stimuli. Nevertheless, the chemosensitivies of these two groups were not identical. NaCl evoked a mean net response about 10X greater in SST_neg_ neurons and BIT elicited a response about 1.5X as great in SST_pos_ cells. However, firing rates were very heterogeneous across neurons and performing an ANOVA on the net firing rates yielded no main effect of neuron type or stimulus and no interaction between these variables (all P’s > .1; mixed ANOVA). However, when we split neurons into groups based on their optimal responsiveness to different stimulus categories (sweet/umami, salt, sour, bitter), clearer differences emerged. A majority (8/14, 57%) of SST_pos_ neurons responded optimally to the bitter stimulus, while none responded best to NaCl (Figure 4). In contrast, 29% of the SST_neg_ neurons were NaCl-best, but only 14% were bitter-best (X^2^, *p* = .023). Indeed, when responses for each neuron were expressed relative to their maximum response (Figure 5), there was a significant interaction between stimulus and SST type (ANOVA) and post-hoc T-tests yielded a significant difference for the BIT response and one that approached significance for NaCl.

**Figure 4.**
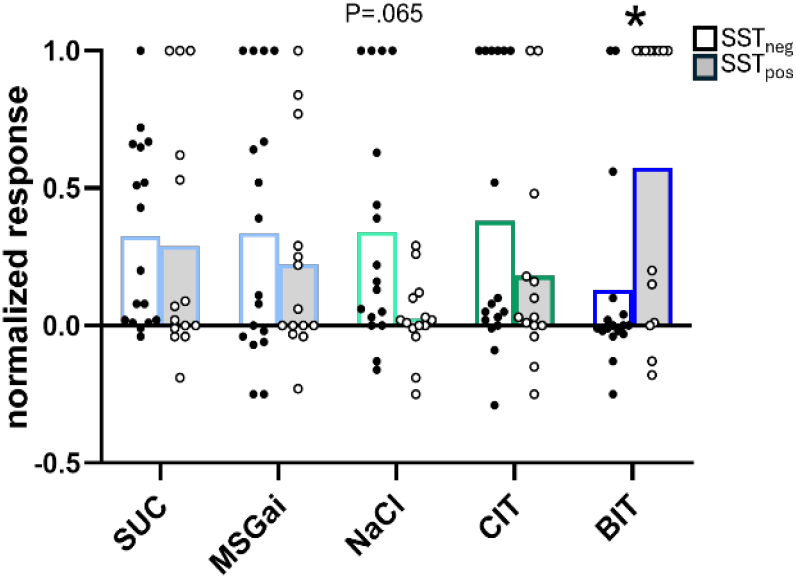
**A**. Heat map for individual neurons showing response profiles normalized to the maximum response. Cells are sorted by their “best” stimulus. **B**. Proportions of SST_neg_ (top) and SST_pos_ (bottom) neurons responding maximally to the different stimulus categories. Color coding in the pie chart matches the colors of the labels for the heat map.

**Figure 5.**
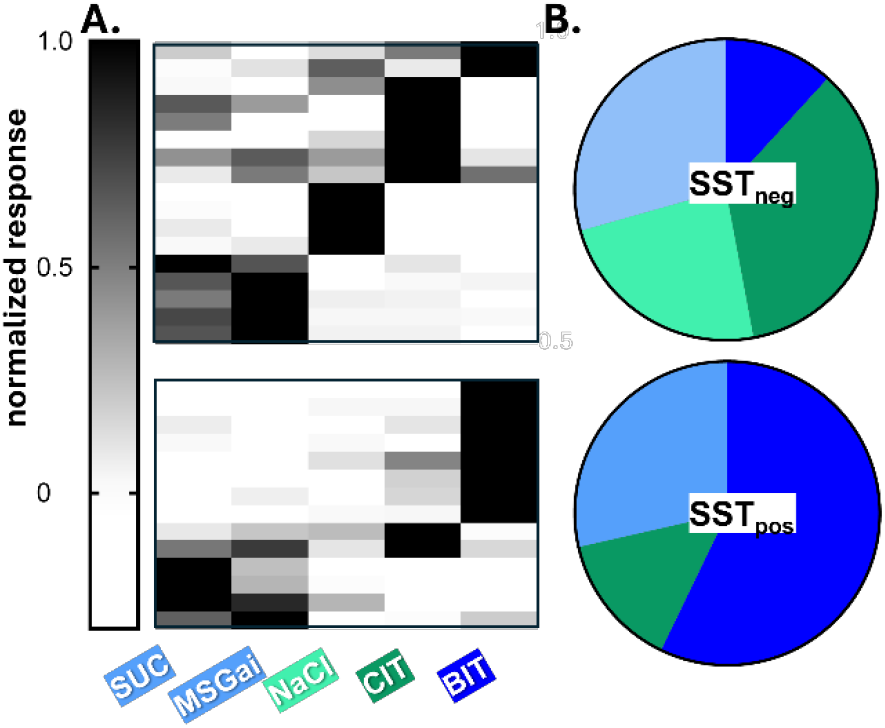
Mean and individual responses to each of the five stimuli normalized to the maximal response. Open bars: SST_neg_; grey shaded bars: SST_pos_ neurons. ANOVA (cell type-*p* = .284, stimulus-*p* =.664, stimulus X cell type-*p* =.012). Bonferroni-adjusted paired t-tests yielded a significant difference for BIT (*p* = .045) and a marginal difference for NaCl (*p* = .065).

### Effect of optogenetic activation of SST circuitry on SST-neurons

In these same mice, we evaluated the effect of light stimulation through the optrode on taste responses of putative SST_neg_ neurons. We presume that this manipulation activated nearby SST_pos_ neurons and fibers, arising from local connections in the NST as well as SST projections from other sources such as the central nucleus of the amygdala (Bartonjo and Lundy 2020, Jin, Fishman et al. 2021) Twelve of 17 SST_neg_ neurons were tested with and without concurrent light stimulation using all five stimuli. Across these 12 cells, the mean response to each stimulus was nominally smaller during light stimulation, but this suppression did not reach statistical significance. However, we were able to test the most effective stimulus for a given neuron with and without concurrent light stimulation for all 17 cells (Figure 6). For the optimal stimulus for each neuron, the mean net response was significantly depressed, with individual neurons consistently responding with a similar or depressed magnitude.

**Figure 6.**
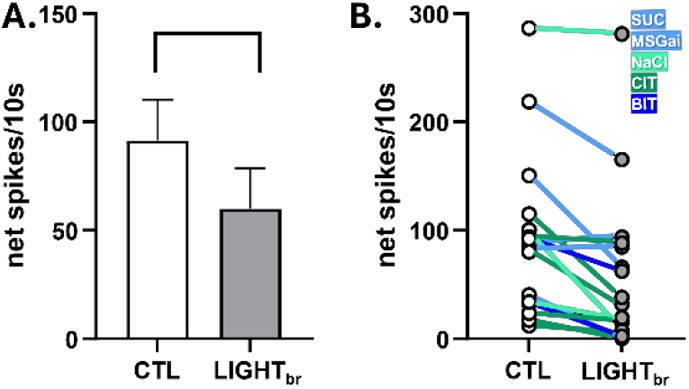
**A**. Responses of SST_neg_ neurons to LIGHT_br_. The mean net response to the “best” stimulus for a given cell was significantly smaller during concurrent stimulation through the optrode (LIGHT_br_) than under control conditions (CTL, paired T-test, *p* =.001). **B**. Responses of individual neurons (N=17). The connecting lines are color-coded according to stimulus. Note that all responses to all qualities could be suppressed by LIGHT_br_.

### Behavioral Studies

Figure 7 shows photomicrographs of injection sites from representative mice in the G_i_ (A) and mC (B) groups; histological results were similar for the other cases. Fluorescence from the injections was centered in rNST with some spread into the overlying vestibular nucleus and underlying dorsal parvocellular reticular formation. Viral label was most robust at a middle level of the rostral zone (i.e., midway between where the NST moves lateral to the IVth ventricle and the rostral limit of the nucleus) but spread throughout the rNST. Minimal label was present caudal to rNST, mainly consisting of fibers and a few cells in the intermediate NST.

**Figure 7.**
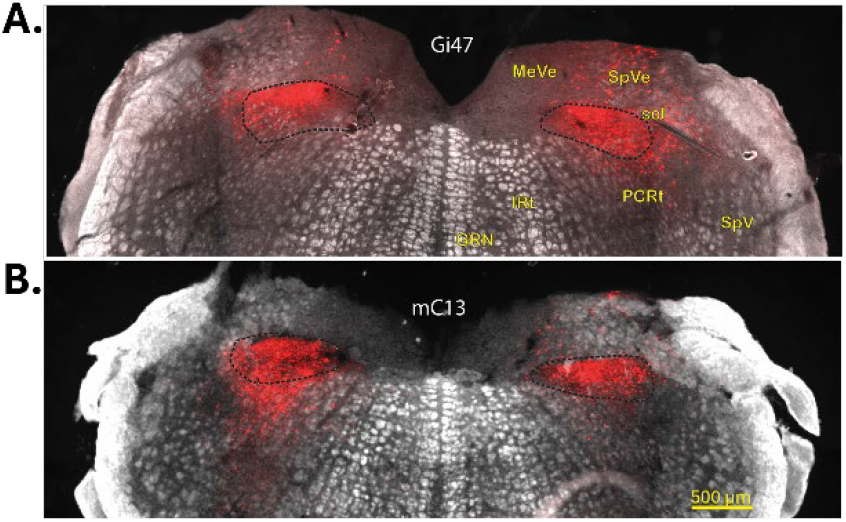
Representative rNST injections of. **A.** pAAV2-hSyn-DIO-hM4D-mCherry virus (Gi47) and **B**. the pAAV2-hSyn-DIO-mCherry (mC13) virus. Fluorescence and darkfield images are superimposed. Black dotted lines show the approximate borders of the nucleus. Abbreviations: GRN-gigantocellular reticular nucleus, Irt-intermediate reticular nucleus, MeVe: medial vestibular nucleus, PCRt-parvicelluar reticular nucleus, sol-solitary tract, SpVe-spinal vestibular nucleus, SpV-spinal trigeminal nucleus.

Inhibiting rNST SST neuron activity had effects on both quinine (QUI) and sucrose (SUC) licking in the brief-access test (Figure 8). In the G_i_ group, compared to I.P. saline injections, mice licked significantly more to QUI at the two highest concentrations, suggesting depressed avoidance, with ANOVA yielding main effects of drug and concentration and an interaction between these variables. In the mC group, only concentration yielded a significant effect. In addition, CNO nominally increased licking for SUC in the 30-300 mM range and ANOVA showed a drug X concentration interaction. In contrast to the effect on QUI licking, increased licking of this preferred stimulus suggests enhanced short-term preference (Figure 8). It is important to note that, although mice licked more for both stimuli at certain concentrations, motivational or motor factors are unlikely factors as neither the number of trials nor water licks were affected by inhibiting SST neural activity (Figure 9). However, the ILI for SUC following CNO was slightly albeit significantly shorter than after saline, consistent with increased licking for SUC in this group. None of these variables were affected in the mC group.

**Figure 8.**
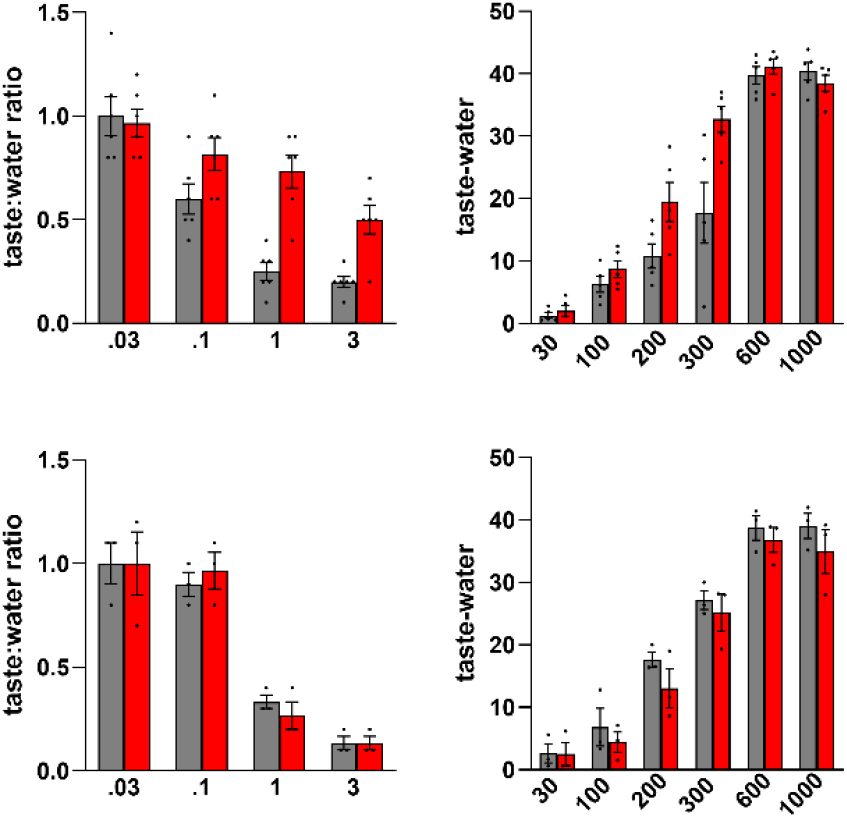
Inhibiting SST neurons suppressed QUI avoidance but augmented SUC preference. **A.** Mice in the Gi group (N = 6) licked more to aversive QUI after CNO than saline injections (ANOVA: drug - *p* = .001, concentration – *p* = 5.29 X E-9; drug X concentration – *p* = .029). **B**. At low to mid-range concentrations, mice in the Gi group (N = 5) licked nominally more SUC after CNO than after saline injections (ANOVA: drug - *p*=.087, concentration – *p* = 9.88 X E-16, drug X concentration – *p* = .006). **C**. Only concentration yielded a main effect for QUI in the mC group (N=3 - Drug-*p* =.777, concentration-*p* = 3.42 e-5, drug X concentration-*p* = .171). **D**. Only concentration yielded a main effect for SUC in the mC group (N = 3), though there was a marginal decrease in licking after CNO across concentrations (N=3, Drug-*p* = 0.054, concentration-*p* =1.5 e-6, drug X concentration-*p* = .609). “*” indicates significant differences based on Bonferroni-adjusted paired T-tests.

**Figure 9.**
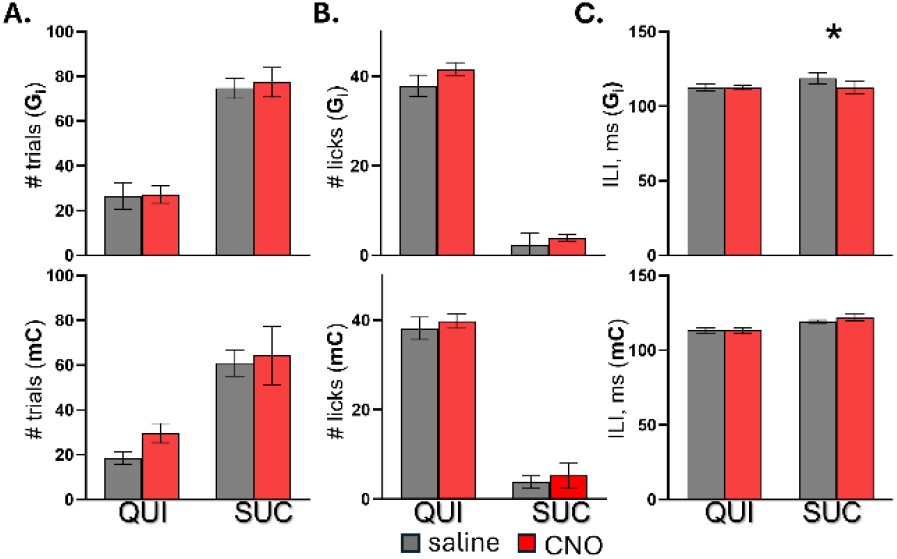
Suppressing neural activity in SST neurons did not generally influence motivational or motor variables. Paired T-tests showed no significant differences between the number of trials **(A)** following saline versus CNO injections for either QUI (*p* = .796 Gi; *p* = .193, mC) or SUC (*p* = .569 Gi; *p* = .698 mC). Likewise, the number of water licks **(B)** was not different for either stimulus or group (*p* = .307, Gi, QUI; *p* = .671, mC, QUI; *p*= .132 Gi, SUC; *p* = .355 mC, SUC). The ILI (C) was not different for qui for either group (*p* = .937, Gi; 1.00, mC or SUC for mC mice (*p* = .344). However, the ILI was slightly though significantly higher after CNO than saline for the Gi group (*p* = .007).

## DISCUSSION

### Somatostatin neurons

In situ hybridization showed that rNST SST neurons are a varied population that are either excitatory or inhibitory. The proportions of both types are substantial, but inhibitory cells are ∼1.7X more prominent. These proportions of glutamatergic and GABAergic SST neurons in rNST are almost identical to reports in the caudal visceral NST (Thek, Ong et al. 2019). Significantly, the existence of sizeable populations of excitatory and inhibitory SST neurons in the nucleus of the solitary tract contrasts with other central regions; for example, SST neurons are largely inhibitory in the cortex and hippocampus (Urban-Ciecko and Barth 2016), and excitatory in the spinal cord (Duan, Cheng et al. 2014).The existence of both excitatory and inhibitory SST neurons point to multiple functional roles for these neurons. We observed a degree of topography in the locations of excitatory and inhibitory SST cells, with the ventral and lateral regions of the nucleus containing a denser collection of VGAT/SST relative to VGLUT2/SST neurons. The significance of this differential distribution is unclear but previous studies suggest that rNST neurons giving rise to projections to the reticular formation are more prominent ventrally and laterally, whereas those projecting to the parabrachial nucleus are most dense dorsally and medially (Travers 1988, Halsell, Travers et al. 1996, Ganchrow, Ganchrow et al. 2014) (reviewed in (Travers and Travers 2018)). Thus, it is possible that SST/VGLUT2 neurons in the dorsal and medial regions are likely to participate in sending excitatory signals in the ascending gustatory pathway, a hypothesis consistent with earlier studies showing that most rNST-PBN projection neurons are glutamatergic (Gill, Madden et al. 1999). In contrast, SST/VGAT neurons in the ventral and lateral regions may be particularly important in modulating taste-elicited oromotor behaviors. Since many GABA NST neurons are interneurons (Davis 1988, Lasiter and Kachele 1988), this modulation may occur largely via intrasolitary interactions. However, direct influences on the reticular formation are also a possibility, since a recent study demonstrated direct projections of GABA neurons to regions outside NST (Shi, Ding et al. 2021).

### Chemosensitivity & SST phenotype

Recordings from rNST neurons optotagged as SST_pos_ or SST_neg_ revealed a more nuanced representation of taste quality by these cell groups than recently proposed (Jin, Fishman et al. 2021). Nevertheless, similar to these earlier findings, many SST_pos_ neurons responded selectively and robustly to a bitter stimulus, QUI+CYC. Moreover, the mean normalized bitter response was larger in SST_pos_ than in SST_neg_ cells and a higher proportion of SST_pos_ neurons responded best to QUI+CYC compared to SST_neg_ neurons. On the other hand, there were also SST_pos_ neurons that responded notably, sometimes optimally, to stimuli evoking other qualities (Figure 5). In fact, except for NaCl, which elicited an above-threshold response in only one SST_pos_ neuron, 29-36% of SST_pos_ neurons responded at above criterion rates of firing to SUC or MSG_ai_, and CIT. Although not commented on in the earlier paper, fiber photometry likewise suggested that SST_pos_ neurons responded to citric acid, sucrose, AceK and an MPG+IMP mixture, though weakly compared to quinine and cycloheximide (see figure 2 in (Jin, Fishman et al. 2021).In agreement with our results, the fiber photometry results also indicated that SST neurons responded minimally if at all to NaCl. However, in apparent contrast to the earlier study, the current observations revealed that, although SST_pos_ neurons responded robustly to the bitter stimulus, 18% of SST_neg_ neurons also exhibited bitter-evoked responses, including two cells that responded optimally to this quality. In sum, we conclude that SST_pos_ neurons are not exclusively responsive to bitter tastants nor are bitter responses restricted to this phenotype. Nevertheless, taste qualities are not represented randomly in SST_pos_ and SST_neg_ n, a finding that supports the existence of definable chemosensitive neuron types. In future studies, it would be interesting to know whether differential chemosensitivity extends to excitatory and inhibitory subtypes of SST_pos_ neurons.

### Modulation of taste responses by activation of SST neurons

When taste responses in SST_neg_ neurons were evaluated in the presence and absence of light_br_ stimulation, an overall suppressive effect was observed. This dampening of taste-evoked activity is likely to have multiple sources, including activating local SST inhibitory neurons and SST inhibitory axons projecting from locations outside NST. One major extrinsic SST input arises from the central nucleus of the amygdala (Saha, Henderson et al. 2002, Bartonjo and Lundy 2020, Jin, Fishman et al. 2021) and multiple studies have demonstrated that most of these neurons are GABAergic (Bartonjo and Lundy 2020). It is worth noting that, although about 36% of SST neurons in the rNST are excitatory, we never observed enhancement of taste responses in SST_neg_ neurons during light_br_ stimulation. This is consistent with what Thek and colleagues (Thek, Ong et al. 2019) observed in the caudal NST in a slice preparation, also using an SST/ChR2 mouse line. They reported that afferent responses evoked by solitary tract stimulation were often reduced by concurrent stimulation with light. However, they did not observe any increases. Moreover, they noted that light stimulation evoked IPSPs in 81% of neurons but EPSPs in just 4%. Although the remaining neurons responded with a combined IPSP/EPSP response, IPSPs were monosynaptic and EPSP’s were polysynaptic and longer-latency. These authors concluded that the SST/VGLUT2 neurons in the caudal NST likely projected outside the nucleus, rather than serving as modulatory interneurons. This seems likely for the rNST as well.

### Function

The fact that rNST SST neurons are comprised of both excitatory and inhibitory populations was reflected in the behavioral results. When the activity of SST neurons was suppressed with DREADDS, we observed increased licking to quinine, implying that the intensity of this aversive stimulus was dampened. These behavioral results agree with those of (Jin, Fishman et al. 2021), who likewise reported increased licking to quinine in 15-minute tests after ablation of SST neurons using diphtheria toxin (DTA). These behavioral findings are consistent with the current neurophysiological results which revealed a large population of bitter-responsive SST neurons. It is also consistent with the robust bitter responses in SST neurons revealed by fiber photometry (Jin, Fishman et al. 2021). Because neurons expressing glutamate are the main rNST population that projects to targets outside the nucleus (Gill, Madden et al. 1999), it seems most parsimonious that this behavioral effect on bitter aversion is due to suppressing the activity of glutamatergic bitter-responsive SST cells.

Suppressing SST neurons with DREADDS did not dampen licking to the preferred stimulus sucrose, the expected effect if this manipulation primarily silenced excitatory sucrose-responsive neurons. Instead, increased licking was observed. In contrast, (Jin, Fishman et al. 2021) reported no effect on licking to a different sweet stimulus, AceK, after DTA of SST rNST neurons. This discrepancy may reflect methodology since the other group used only one concentration of the sweet stimulus and compared licking between groups, whereas the current study used a more sensitive within-subject comparison and multiple sucrose concentrations. Interestingly, the behavioral enhancement to sucrose occurred despite neurophysiological evidence that some, albeit a smaller population of SST_pos_ neurons, are optimally responsive to sucrose. We speculate that this reflects a dominant effect of removing inhibitory influences of SST/VGAT neurons, i.e. disinhibition, instead of suppressing those SST neurons that express VGLUT2. However, because we did not manipulate SST/VGAT and SST/VGLUT2 neurons independently, conclusions about the underlying circuitry remain tentative. It would be interesting to use intersectional genetics to target these populations separately to untangle the effects.

### Concluding Remarks

The current study presents three lines of evidence demonstrating heterogeneous functions of rNST SST neurons. These cells are a mixture of excitatory and inhibitory phenotypes, respond to tastants representing multiple qualities, and have influences on behavioral responses to both sweet and bitter stimuli. Nevertheless, the gustatory function of these neurons is not indiscriminate. SST neurons have a robust, dominant sensitivity to bitter stimuli relative to other qualities. Moreover, lack of activity in these cells notably suppresses responding to bitter, but not sweet stimuli. Overall, these results support a functional segregation of cell types at the initial stage of central gustatory processing, but at the same time demonstrate that it is not absolute. This conclusion is consistent with previous data gathered without the benefit of genetically defined cell types (Spector and Travers 2005, Ohla, Yoshida et al. 2019, Travers and Spector 2021) and supports a more nuanced scheme for how individual cells represent gustatory quality.

## Acknowledgments

This work was supported by NIH DC06112 to SPT. We are grateful to Emma Gutarts and Sidney Li for meticulous cell counts and technical contributions. Drs. Joseph B. Travers and Alan C. Spector made valuable comments on the manuscript.

## References

Ascencio Gutierrez, V., L. E. Martin, A. Simental-Ramos, K. F. James, K. F. Medler, L. A. Schier and A. M. Torregrossa (2024). “TRPM4 and PLCbeta3 contribute to normal behavioral responses to an array of sweeteners and carbohydrates but PLCbeta3 is not needed for taste-driven licking for glucose.” Chem Senses 49.

Bartonjo, J. J. and R. F. Lundy (2020). “Distinct Populations of Amygdala Somatostatin-Expressing Neurons Project to the Nucleus of the Solitary Tract and Parabrachial Nucleus.” Chem Senses 45(8): 687–698.

Bellusci, L., S. N. Garcia DuBar, M. Kuah, D. Castellano, V. Muralidaran, E. Jones, A. M. Rozeboom, R. A. Gillis, S. Vicini and N. Sahibzada (2022). “Interactions between Brainstem Neurons That Regulate the Motility to the Stomach.” J Neurosci 42(26): 5212–5228.

Boxwell, A. J., Y. Yanagawa, S. P. Travers and J. B. Travers (2013). “The mu-opioid receptor agonist DAMGO presynaptically suppresses solitary tract-evoked input to neurons in the rostral solitary nucleus.” J Neurophysiol 109(11): 2815–2826.

Breza, J. M. and S. P. Travers (2016). “P2×2 Receptor Terminal Field Demarcates a “Transition Zone” for Gustatory and Mechanosensory Processing in the Mouse Nucleus Tractus Solitarius.” Chem Senses 41(6): 515–524.

Davis, B. J. (1988). “Computer-generated rotation analyses reveal a key three-dimensional feature of the nucleus of the solitary tract.” Brain Res Bull 20(5): 545–548.

Deng, H., X. Xiao, T. Yang, K. Ritola, A. Hantman, Y. Li, Z. J. Huang and B. Li (2021). “A genetically defined insula-brainstem circuit selectively controls motivational vigor.” Cell 184(26): 6344–6360 e6318.

Duan, B., L. Cheng, S. Bourane, O. Britz, C. Padilla, L. Garcia-Campmany, M. Krashes, W. Knowlton, T. Velasquez, X. Ren, S. Ross, B. B. Lowell, Y. Wang, M. Goulding and Q. Ma (2014). “Identification of spinal circuits transmitting and gating mechanical pain.” Cell 159(6): 1417–1432.

Ganchrow, D., J. R. Ganchrow, V. Cicchini, D. L. Bartel, D. Kaufman, D. Girard and M. C. Whitehead (2014). “Nucleus of the solitary tract in the C57BL/6J mouse: Subnuclear parcellation, chorda tympani nerve projections, and brainstem connections.” J Comp Neurol 522(7): 1565–1596.

Geran, L. C. and S. P. Travers (2006). “Single neurons in the nucleus of the solitary tract respond selectively to bitter taste stimuli.” J Neurophysiol 96(5): 2513–2527.

Gill, C. F., J. M. Madden, B. P. Roberts, L. D. Evans and M. S. King (1999). “A subpopulation of neurons in the rat rostral nucleus of the solitary tract that project to the parabrachial nucleus express glutamate-like immunoreactivity.” Brain Res 821(2): 251–262.

Giza, B. K. and T. R. Scott (1983). “Blood glucose selectively affects taste-evoked activity in rat nucleus tractus solitarius.” Physiol Behav 31(5): 643–650.

Halsell, C. B., S. P. Travers and J. B. Travers (1996). “Ascending and descending projections from the rostral nucleus of the solitary tract originate from separate neuronal populations.” Neuroscience 72(1): 185–197.

Inui-Yamamoto, C., G. D. Blonde, F. Schmid, L. Mariotti, M. Campora, T. Inui, L. A. Schier and A. C. Spector (2020). “Neural Isolation of the Olfactory Bulbs Severely Impairs Taste-Guided Behavior to Normally Preferred, But Not Avoided, Stimuli.” eNeuro 7(2).

Jin, H., Z. H. Fishman, M. Ye, L. Wang and C. S. Zuker (2021). “Top-Down Control of Sweet and Bitter Taste in the Mammalian Brain.” Cell 184(1): 257–271 e216.

Kalyanasundar, B., G. D. Blonde, A. C. Spector and S. P. Travers (2020). “Electrophysiological responses to sugars and amino acids in the nucleus of the solitary tract of type 1 taste receptor double-knockout mice.” J Neurophysiol 123(2): 843–859.

Kalyanasundar, B., G. D. Blonde, A. C. Spector and S. P. Travers (2023). “A Novel Mechanism for T1R-Independent Taste Responses to Concentrated Sugars.” J Neurosci 43(6): 965–978.

Kalyanasundar, B., A. Harley, C. Klimovich and S. Travers (2025). “Chemogenetic suppression of NST GABA neurons reveals inhibition of behavioral responses to sucrose and quinine.” Physiol Behav 295: 114889.

Kalyanasundar, B., C. Klimovich, S. Li, E. Gutarts and S. Travers (2022). “Chemogenetic Inhibition Of Somatostatin Neurons In The Nucleus Of The Solitary Tract Differentially Modulates Bitter And Sweet Taste Signals.” 44th Annual Meeting of Association for Chemoreception Sciences Annual Meeting Abstracts: #315.

King, C. T., S. P. Travers, N. E. Rowland, M. Garcea and A. C. Spector (1999). “Glossopharyngeal nerve transection eliminates quinine-stimulated fos-like immunoreactivity in the nucleus of the solitary tract: implications for a functional topography of gustatory nerve input in rats.” J Neurosci 19(8): 3107–3121.

Lasiter, P. S. and D. L. Kachele (1988). “Organization of GABA and GABA-transaminase containing neurons in the gustatory zone of the nucleus of the solitary tract.” Brain Res Bull 21(4): 623–636.

Lemon, C. H. (2017). “Modulation of taste processing by temperature.” Am J Physiol Regul Integr Comp Physiol 313(4): R305–R321.

Mai, L., A. S. Ramos, A. H. Jung, D. de Monteiro, I. Vu, T. Saputera, J. Fan, T. Adkins, D. Dickerson, D. W. Pittman, S. Chometton and L. A. Schier (2025). “Nutritional regulation of metabolism-dependent and-independent glucosensing in the mammalian taste system.” Mol Metab 103: 102280.

McLean, I. and P. Nakane (1974). “Periodate-lysine-paraformaldehyde fixative. A new fixation for immunoelectron microscopy.” J Histochem Cytochem 12: 1077–1083.

Nakajima, K. I. (2024). “Recent advances in the characterization of genetically defined neurons that regulate internal-state-dependent taste modification in mice.” Physiol Rep 12(21): e70106.

Nakamura, K. and R. Norgren (1995). “Sodium-deficient diet reduces gustatory activity in the nucleus of the solitary tract of behaving rats.” Am J Physiol 269(3 Pt 2): R647–661.

Ohla, K., R. Yoshida, S. D. Roper, P. M. Di Lorenzo, J. D. Victor, J. D. Boughter, M. Fletcher, D. B. Katz and N. Chaudhari (2019). “Recognizing Taste: Coding Patterns Along the Neural Axis in Mammals.” Chem Senses 44(4): 237–247.

Pilato, S. A., F. P. O’Connell, J. D. Victor and P. M. Di Lorenzo (2024). “Electrophysiological responses to appetitive and consummatory behavior in the rostral nucleus tractus solitarius in awake, unrestrained rats.” Front Integr Neurosci 18: 1430950.

Preibisch, S., S. Saalfeld and P. Tomancak (2009). “Globally optimal stitching of tiled 3D microscopic image acquisitions.” Bioinformatics 25(11): 1463–1465.

Saha, S., Z. Henderson and T. F. Batten (2002). “Somatostatin immunoreactivity in axon terminals in rat nucleus tractus solitarii arising from central nucleus of amygdala: coexistence with GABA and postsynaptic expression of sst2A receptor.” J Chem Neuroanat 24(1): 1–13.

Schier, L. A., C. Inui-Yamamoto, G. D. Blonde and A. C. Spector (2019). “T1R2+T1R3-independent chemosensory inputs contributing to behavioral discrimination of sugars in mice.” Am J Physiol Regul Integr Comp Physiol 316(5): R448–R462.

Shi, M. Y., L. F. Ding, Y. H. Guo, Y. X. Cheng, G. Q. Bi and P. M. Lau (2021). “Long-range GABAergic projections from the nucleus of the solitary tract.” Mol Brain 14(1): 38.

Spector, A. C. and S. P. Travers (2005). “The representation of taste quality in the mammalian nervous system.” Behav Cogn Neurosci Rev 4(3): 143–191.

Thek, K. R., S. J. M. Ong, D. C. Carter, J. K. Bassi, A. M. Allen and S. J. McDougall (2019). “Extensive Inhibitory Gating of Viscerosensory Signals by a Sparse Network of Somatostatin Neurons.” J Neurosci 39(41): 8038–8050.

Travers, J. B. (1988). “Efferent projections from the anterior nucleus of the solitary tract of the hamster.” Brain Res 457(1): 1–11.

Travers, J. B., K. Herman, J. Yoo and S. P. Travers (2007). “Taste reactivity and Fos expression in GAD1-EGFP transgenic mice.” Chem Senses 32(2): 129–137.

Travers, J. B. and S. P. Travers (2018). Microcircuitry of the rostral nucleus of the solitary tract. Handbook of Brain Microcircuits G. M. Shepherd and S. Grillner, Oxford.

Travers, S. P., B. Kalyanasundar, J. Breza, G. Houser, C. Klimovich and J. Travers (2022). “Characteristics and Impact of the rNST GABA Network on Neural and Behavioral Taste Responses.” eNeuro 9(5).

Travers, S. P. and R. Norgren (1995). “Organization of orosensory responses in the nucleus of the solitary tract of rat.” J Neurophysiol 73(6): 2144–2162.

Travers, S. P. and A. C. Spector (2021). Anatomical Organization and Coding in the Gustatory System: A Functional Perspective. Oxford Research Encylopedias. Neuroscience.

Urban-Ciecko, J. and A. L. Barth (2016). “Somatostatin-expressing neurons in cortical networks.” Nat Rev Neurosci 17(7): 401–409.

Urban, D. J. and B. L. Roth (2015). “DREADDs (designer receptors exclusively activated by designer drugs): chemogenetic tools with therapeutic utility.” Annu Rev Pharmacol Toxicol 55: 399–417.

Wang, M. and R. M. Bradley (2010). “Properties of GABAergic neurons in the rostral solitary tract nucleus in mice.” J Neurophysiol 103(6): 3205–3218.

Whitehead, M. C. (1988). “Neuronal architecture of the nucleus of the solitary tract in the hamster.” J Comp Neurol 276(4): 547–572.

Wu, A., G. Dvoryanchikov, E. Pereira, N. Chaudhari and S. D. Roper (2015). “Breadth of tuning in taste afferent neurons varies with stimulus strength.” Nat Commun 6: 8171.

Zhang, J., H. Jin, W. Zhang, C. Ding, S. O’Keeffe, M. Ye and C. S. Zuker (2019). “Sour Sensing from the Tongue to the Brain.” Cell 179(2): 392–402 e315.

